# Simple attributes predict the importance of plants as hosts to the richness of fungi and arthropods

**DOI:** 10.1101/2020.04.17.046292

**Authors:** Hans Henrik Bruun, Ane Kirstine Brunbjerg, Lars Dalby, Camilla Fløjgaard, Tobias G. Frøslev, Simon Haarder, Jacob Heilmann-Clausen, Toke T. Høye, Thomas Læssøe, Rasmus Ejrnæs

## Abstract

Consumers constitute the vast majority of global terrestrial biodiversity. Yet, local consumer richness is poorly understood. Plant species richness offers a simple hypothesis to how the diversification of carbon substrates may promote the diversity of arthropods and fungi. We took this one step further and used databases on plant-consumer interaction links to derive the richness of associated biota per plant species (link score). Using a species inventory of 130 sites we investigated 1) how well the link score could be predicted by plant attributes and 2) if the sum of plant species’ observed or predicted link scores could predict site richness of arthropods and macrofungi better than plant species richness alone. We found plant link scores to be positively related to plant size, abundance, nativeness and ectomycorrhizal status. Link based indices generally improved prediction of richness, stressing the importance of plants as niche space for the megadiverse groups of insects and fungi.

## Introduction

Plants constitute an important part of biodiversity in their own right, but in addition provide resource and habitat to all terrestrial heterotrophic biodiversity, including the megadiverse groups of arthropods and fungi. The link from plant diversity to consumer diversity is modulated by the degree of host specialism among consumers. Most species of insects, mites and fungi associated with living plants are strongly specialised, i.e. dependent on a single or a few plant species as both resource and habitat (Strong, Lawton & Southwood 1984; Hawksworth 2001). Even many decomposers show a high degree of specialism, due to after-life effects of plant structure and chemical composition (Heilmann-Clausen *et al.* 2016). Generalist arthropod and fungal species are relatively few in number. Cascading effects from plants to the third and fourth trophic levels have been demonstrated and may be particular important to specialist parasitoids (Godfray 1995).

The direct effect of plants on the trophic levels above is encapsulated in the *ecospace* dimension coined *expansion* signifying the build-up and diversification of organic carbon in the ecosystem (Brunbjerg *et al.* 2017b). However, empirical predictive power of plants on consumer richness may result from both direct effects of plant diversity or from indirect effects of consumers and producers responding in similar ways to extrinsic factors (Kemp & Ellis 2017), in particular abiotic properties such as microclimate, soil moisture, soil nutrients and inorganic substrates, i.e. the position component of ecospace (Brunbjerg *et al.* 2017b). Thus, testing the effect of plant diversity on consumer diversity must account also for the effects of the abiotic environment.

Prediction of local site biodiversity provides essential knowledge to prioritization of conservation efforts, apart from being an intriguing task in itself. While biodiversity surrogacy has been questioned in general (Larsen, Bladt & Rahbek 2009; Lindenmayer *et al.* 2015), plant species richness has proven to promote multi-taxon diversity, provided that – on top of plant species richness – plant species identities are used for bioindication of key habitat conditions (Brunbjerg *et al.* 2018). However, it remains to be tested if higher predictive power may be attained after including plant species’ value as food and habitat to consumer species.

The question goes deeper than surrogacy, however. Some plant species support a much greater diversity of associated consumers than others. So plant identity may be more important than plant species number to local consumer diversity. There are however only few regions in the world where the consumer to host links have been mapped adequately, and it is therefore interesting to know if host attractiveness can be predicted.

A number of key plant attributes are known to be related to the richness of associated arthropod and fungal species. In particular properties revolving around plant apparency and predictability as a resource have been found important (Feeny 1976), i.e. species range size and local abundance, body size and life span, time since immigration and nativeness (Southwood 1961; Lawton & Schröder 1977; Kennedy & Southwood 1984; Brändle & Brandl 2001; Miller 2012). In addition, chemical defences and phylogenetic isolation are among the proposed plant determinants of arthropod richness (Tahvanainen & Niemelä 1987; Brändle & Brandl 2006) and, for symbiotic fungi, plant species’ ability to form mycorrhiza of different types (Tedersoo *et al.* 2015). The literature regarding arthropods is much bigger than for fungi; few analyses have combined the two (Strong & Levin 1979; Brändle & Brandl 2003), and even fewer have taken investigations from the level of whole biotas to the level of local communities.

Here we use species richness of vascular plants, arthropods and macrofungi surveyed at 130 sites representing all terrestrial ecosystems in a region as a study case. We used independently recorded plant-consumer interaction links to derive the size of the associated consumer biota per plant species (observed link score). This observed link score was modelled from plant attributes, such as range size and growth form (predicted link score). Link scores were summed over the plant species occurring in each site to obtain an observed and predicted link sum per site. In addition to modelling total fungal and arthropod richness over study sites, the funga and arthopod fauna were divided into functional response groups, which we expected were controlled by different abiotic and biotic drivers.

Specifically we asked: 1) Which plant attributes can predict the potential number of interaction links between plants and associated arthropods and fungi, 2) Can link sum predict the observed species richness of fungi and arthropods (and functional subgroups thereof) on the scale of communities, better than raw plant species richness? 3) Can the observed link score be substituted by a trait-predicted link sum in the prediction of insect and fungi richness?

## Methods

### Study area and collection of biodiversity data

The study area was Denmark (Fig. 1). In the field, we collected data from 130 sites, each with an area of 40 × 40 m, deemed to be homogenous with respect to topography and vegetation structure, but accepting the inherent heterogeneity of some habitat types. The study design aimed at coverage of the major environmental gradients, including naturalness of habitat (i.e. the intensity of silviculture and agriculture). Thirty sites were allocated to cultivated habitats and 100 sites to natural habitats. The cultivated subset was stratified according to major land use classes and the natural subset was stratified across gradients in soil fertility, soil moisture and vegetation openness. The design has been described in detail by Brunbjerg et al. (2017a).

**Fig. 1.**
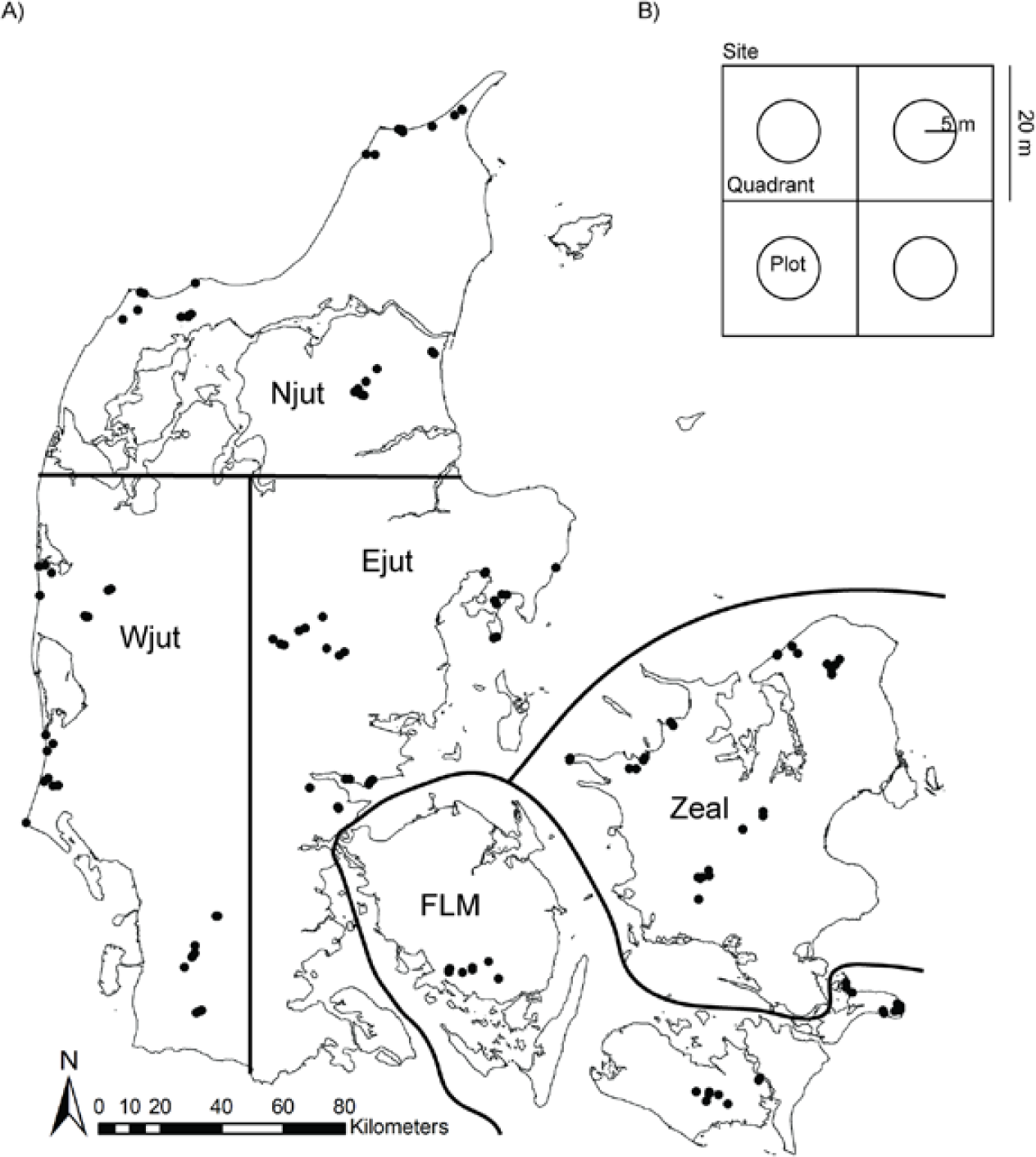
Map of Denmark showing the location of the 130 sites grouped into 15 clusters within five regions (Njut: Northern Jutland, Wjut: Western Jutland, Ejut: Eastern Jutland, FLM: Funen, Lolland, Møn, Zeal: Zealand). B) Site layout with four 20 × 20 m quadrants each containing a central 5 m radius circular plot.

We collected data on the occurrence of vascular plants, macrofungi and arthropods, aiming for an unbiased and representative assessment of the multi-taxon species diversity in each of the 130 sites. For vascular plants, the sampling included abundance assessment on a coarse scale. Each site was divided into four quadrants and, at the centre of each quadrant, a 50 × 50 cm inner quadrat embedded in circular plot (5 m radius) was situated. Presence of species was recorded in inner quadrats and 5 m plots separately, in addition to records for the whole site. Plant species judged by visual inspection to be dominant at site-level were noted. For the present analysis, we assigned an ordinal abundance of 3 to plants species either judged as dominants or recorded in all four inner quadrats. Plant species recorded in at least one inner quadrat and at least one 5 m plot were assigned an ordinal abundance of 2. The remaining species were assigned a weight of 1. Proxy variables for site environmental conditions were derived from site plant lists by bioindication using Ellenberg Indicator Values for light, soil nutrients, soil moisture and soil pH (Ellenberg *et al.* 1991; Brunbjerg *et al.* 2018).

The data for vascular plants may be considered as good as exhaustive, while for the remaining species groups, which are more demanding to find or catch and to identify, the data represent a reproducible and unbiased sampling effort across the 130 sites. For arthropods, we operated a standard set of pitfall traps, yellow pan traps and Malaise traps during two set periods in 2014. Furthermore, two active-search approaches were used to retrieve externally and internally plant-feeding arthropods, respectively: 1) Sweep netting and beating with a focus on bugs, cicadas and leaf beetles, 2) strategic search for plant galling and mining arthropods, including non-galling Cecidomyiinae. Metabarcoding was applied to soil samples from all 130 sites in order to obtain OTU-richness estimates for cryptic soil biota (see Brunbjerg *et al.* 2017a for details). OTUs were derived using the nuclear ribosomal ITS2 marker for fungi and the nuclear16S rRNA marker for arthropods. The fungal OTU data were split into Agaricomycetes, which largely overlap with the macrofungi surveyed, and non-Agaricomycetes, which is a large and phylogenetically heterogeneous group that mostly goes undetected in traditional surveys, using the UNITE database (Nilsson *et al.* 2019). For full details on data collection, see Brunbjerg et al. (2017a).

### Interaction data and link score calculation

We extracted data for interaction links from existing databases for each plant species found at the 130 field sites. For both fungi and arthropods, the estimation had to take into account that many interaction links have been recorded at the level of plant genus, either because consumers do not discriminate between different species of the same plant genus (Savile 1979) or because of incomplete identification of the host by the human observer.

We mined a Danish fungal database (https://svampe.databasen.org) for fungi and the Biological Records Centre’s host plant database for arthropods (http://www.brc.ac.uk/dbif/hosts.aspx) for reported links to the list of vascular plants found in the studied sites, including common plant name synonyms. We included interaction links reported at plant subspecies, species or genus levels.

The fungal database is based on repeated field observations of fungi and the arthropod databases is based on unique reported links and we thus had to be handle them in slightly different ways in order to obtain comparable data for analysis. A particular challenging task was to distribute fungal and arthropod links reported at the plant genus level onto the plant species belonging to that genus. The procedure is detailed below.

The Danish fungal database consists of observations of fungi made by citizens and professionals, with records accompanied by observational data on substratum, i.e. live plants or dead plant parts, or – for ectomycorrhizal fungi – close association with plants, identified at least to plant genus.

The British BRC host plant database is a meta-database, compiling arthropod-plant associations reported in the scientific literature. The geographic focus is Great Britain and adjacent continental Europe. The database is regularly updated, but curation of arthropod taxonomy and nomenclature is not better than the most recent source for each arthropod group, which for many little studied groups may be quite old. Because of the vast number of literature sources, cleaning the arthropod names for synonyms was considered intractable. However, despite fair criticisms of biases towards common plant species, the reliability of published host records is very well supported (Brändle & Brandl 2001). We retrieved arthropod links for all vascular plant species found across the 130 field sites, under the assumption that the BRC database would give a fair picture of the size of the total coterie of associated arthropods on Danish plants.

### Plant-associated fungi

We retrieved all records of fungi having at least one reported association with a vascular plant at the species or genus level (n = 255 700). Removing duplicate links and filtering to the total list of vascular plant species found at the 130 study sites led to a reduction of data entries to 20 309 links between 4 549 fungal and 538 vascular plant taxa (at species or genus level). For each fungal species, each of its plant links were given a weight corresponding to 1 divided by the number of linked plant genera for that fungus. Thus, all fungal species would contribute identical total weights to the final index, but a specialist fungus would contribute more to the link score of its host plant than would a generalist fungus. These link weights were summed for each vascular plant species over all fungal taxa at both species and genus level, accepting that some fungal species had links reported at both levels. When calculating the final link score for a plant species, we allocated the plant genus score to the species belonging to a given genus in the following way:

1. Plant species having link records at both the species and genus levels were allocated a percentage of the genus link score proportional to their species-level link score relative to the link score of the species within the genus with the highest species-level link score. This rule applied to 265 plant species.
2. Plant species belonging to a genus with link records at the genus level only (no species-level records for any constituent species) were allocated an arbitrary 90% of the genus link score, equal for all species in the genus. This rule applied to 104 plant species.
3. Plant species having link records at the genus level only, but belonging to a genus containing other species with species-level records, were scored as “NA”, based on the argument that neither zero nor a positive link score would be correct. This applied to 245 plant species.
4. Plant species without reported fungal links at neither genus nor species level were given a zero link score. This applied to 293 plant species.

We calculated the final fungal link score for each plant as the sum of the species score and the share of the genus score.

#### Phytophagous arthropods

For the total list of plant species encountered at the 130 sites, we retrieved all reported interactions involving insects and mites from the BRC host plant database. Interactions reported on plant subspecies level were merged on the parent species level. We found 30 895 interaction links, involving 6 870 arthropod species and 1427 vascular plant taxa (at species or genus level), of which 37 % were reported at the plant genus level. Similarly to the procedure for fungi, each plant link of an arthropod species were given a weight corresponding to 1 divided by the number of linked plant genera. For arthropods, however, we gave priority to plant genus links and only used species-level links in case no links were reported at genus level. The link score at plant genus level was allocated to all constituent species equally. While this may often be correct, it may also sometimes imply unwarranted link points to exotic or biologically deviating members of a plant genus. In order to compensate for this possible bias and give some priority to species-specific links, we decided to assign triple weight to link points reported at species level. The final link score for a plant was calculated as the sum of genus-level link weights and three times the sum of species level link weights.

### Plant attributes

In order to re-cast the interaction link score of plants species in terms of their traits, we compiled plant attributes for the plant species found across the 130 field sites. Information on ectomycorrhiza was extracted from the MycoFlor database (Hempel *et al.* 2013). Information on nativeness of plant species at 1) national scale, 2) at the European scale and 3) nativeness of the genus, were taken from Buchwald *et al.* (2013). Plant species were assigned to one of the following taxonomic groups: Angiosperm, Gymnosperm and Pteridophyte, based on standard plant classification. Lifespan was scored as 1) annuals + short-lived perennials, 2) medium-lived perennials, 3) long-lived perennials. Lifeform was scored as 1) tree = macrophanerophyte, 2) shrub+liana = nanophanerophyte, 3) dwarf-shrub = hemiphanerophyte, 4) herb = hemicryptophyte + geophyte + chamaephyte + therophyte + hydrophyte + pseudophanerophyte (Klotz, Kühn & Durka 2002). Plant body size was based on maximum canopy height and re-classified as 1) huge, 2) large, 3) medium-sized, 4) small and 5) tiny, following the LEDA trait data base (Kleyer *et al.* 2008). These attributes were extracted using the R-package TR8 (Bocci 2015). We used family, genus and species descriptions in Hansen (2004) for filling gaps in height information. Plant species regional occupancy was extracted from a national plant survey (Atlas Flora Danica), carried out in 5 × 5 km grid cells, of which 1300 were thoroughly surveyed, resulting in reliable presence-absence data (Hartvig & Vestergaard 2015). Species incidence frequency across reference grid cells was re-coded as High (> 0.75), Moderate (0.26 – 0.75) and Low (< 0.25) occupancy. Because of a bimodal frequency distribution, there were approximately equal numbers of species in the three occupancy classes.

We modelled plant species link score, for fungi and arthropods separately, in response to plant attributes using a linear modelling approach with ectomycorrhizal status, national nativeness, European nativeness, nativeness on the genus level, taxonomic group, life form, lifespan, body size and regional occupancy as explanatory variables. All explanatory variables were coded as factor variables (nominal), fungi link score was log-transformed and insect link score was square-root transformed. Model performance was assessed with type III sum of squares based on reducing a full model with the least significant variable until all variables were significant. The resulting regression models were used to predict the expected number of fungal and arthropod links per plant species based on species traits. The resulting metric is, henceforth, called ‘predicted link score’ as opposed to the ‘observed link score’ based on databases. The correlation between observed and predicted links scores across species was assessed with Spearman rank correlation.

### Link sum per site

For each of the 130 sites, we calculated a simple sum of link scores as well as a weighted link sum, the latter using plant species abundance as weight. The use of plant abundance as weight was based on the reasoning that the local abundance of a plant species would increase the chance that the plant was used as host by fungal or insect species. Simple and weighted link sums were calculated for both observed and predicted link scores. These link sums for a given site would increase with local plant species richness and with the value of the plant species present to fungal and arthropod associates, and thus could also be seen as a link-weighted plant species richness of the site.

### Testing the prediction of biodiversity by interaction scores

We tested the predictive power of interaction link scores on observed multi-taxon species richness data from the 130 field sites. We used total observed species richness of fungi and arthropods as response variable, but also investigated models for functional subgroups of fungi and arthropods, divided according to their relation to plants as resources and ecospace at large. Fungi were divided into symbionts (mainly ectomycorrhizal fungi, but also including biotrophic parasites) and decomposers (saprotrophs). Arthropods were divided into 1) predators, 2) flyers, 3) externally feeding herbivores and 4) internally feeding arthropods, i.e. gallers and miners. The group ‘flyers’ differ from the trophically defined subgroups and was defined by mode of movement and dispersal, reflecting an assumed decoupling of adult and juvenile life stages.

We modelled species richness with GLM, using negative binomial error structure to account for frequent overdispersion of Poisson models. For each taxonomic response group we made a bivariate GLM in response to the link sum. In order to avoid confounding effects from variation in the abiotic environment potentially co-varying with plant link scores, we subsequently ran parallel GLM modelling, in which community mean Ellenberg Indicator Values for light, soil nutrients, soil moisture and soil pH were added to the models as co-variables. We applied multiple regression to test if link sum remained important after fitting a general environmental calibration of the habitat. Both types of model were made for three different sets of plant host richness variables: 1) simple plant richness (corresponding to a null hypothesis of all plant species having equal abundance and equal value as consumer species’ resource), 2) the observed link sum and 3) the predicted link sum. We log-transformed these plant ecospace variables, as this led to decreasing model AIC in most cases – particularly for response groups with strong dependence on host richness. We also modelled abundance-weighted plant richness, but results were almost identical to the simple richness models, so only the latter will be reported here.

## Results

The plant taxa with most fungal interaction links from our database were *Fagus sylvatica, Quercus robur, Picea abies, Pinus sylvestris* and *Salix cinerea*, i.e. all woody plant, and well aligned with previous syntheses (Heilmann-Clausen *et al.* 2016). The herbaceous plants with most fungal links were *Phragmites australis* and *Carex paniculata*. For arthropods, the plant taxa with most interaction links were *Salix* (e.g. *S. cinerea* and *S. repens*), *Quercus* (*Q. robur*), *Pinus sylvestris, Betula* and *Populus* (*P. tremula*), again woody plants and similarly in agreement with previously published evidence (Kelly & Southwood 1999; Brändle & Brandl 2003). The herbaceous plants with most fungal links were *Achillea millefolium* and *Medicago sativa*.

The trait based model of fungal link scores across plant species revealed that capacity to form ectomycorrhiza, high regional occupancy, nativeness to Europe and intermediate to long lifespan all had strong positive effects on the fungal link score, while small body size and herbaceous life form had negative effects (Table 1). The adjusted model R^2^ was 0.65. The parallel model of arthropod link score showed that ectomycorrhizal capacity, high regional occupancy, dwarf shrub lifeform (as opposed to tree, herb or shrub+liana) had significantly positive effects on the arthropod link score, while small body size, short life span and fern phylogenetic placement had significantly negative effects (Table 2). The adjusted model R^2^ was 0.45, i.e. somewhat lower than for the fungal model. Link scores for fungi and insects were significantly positively correlated (Rho = 0.66, p < 0.0001).

**Table 1.** Regression model of fungal link score per plant taxon as predicted from plant attributes. Overall model adjusted R-squared 0.6459; F-statistic: 52.57 on 11 and 300 DF, p-value << 0.0001

|  | Coefficient | Std. Error | t value | P value |
| --- | --- | --- | --- | --- |
| Intercept | 2.0858 | 0.5330 | 3.914 | 0.000113 *** |
| Ectomycorrhizal | 1.6682 | 0.2256 | 7.394 | 1.43e-12 *** |
| Native: Europe | 0.6850 | 0.2179 | 3.144 | 0.001834 ** |
| Occupancy: Low | -0.7532 | 0.1314 | -5.732 | 2.42e-08 *** |
| Occupancy: Moderate | -0.3661 | 0.1001 | -3.658 | 0.000300 *** |
| Lifespan: Short | -0.3598 | 0.1368 | -2.631 | 0.008957 ** |
| Life form: Herbaceous | -0.6941 | 0.2617 | -2.652 | 0.008426 ** |
| Life form: Shrub+Liana | -0.1338 | 0.3206 | -0.417 | 0.676661 |
| Life form: Tree | 0.5542 | 0.4551 | 1.218 | 0.224311 |
| Body size: Large | -0.2523 | 0.2688 | -0.939 | 0.348619 |
| Body size: Medium | -0.1369 | 0.4280 | -0.320 | 0.749322 |
| Body size: Small | -0.9172 | 0.4362 | -2.103 | 0.036323 * |

**Table 2.**
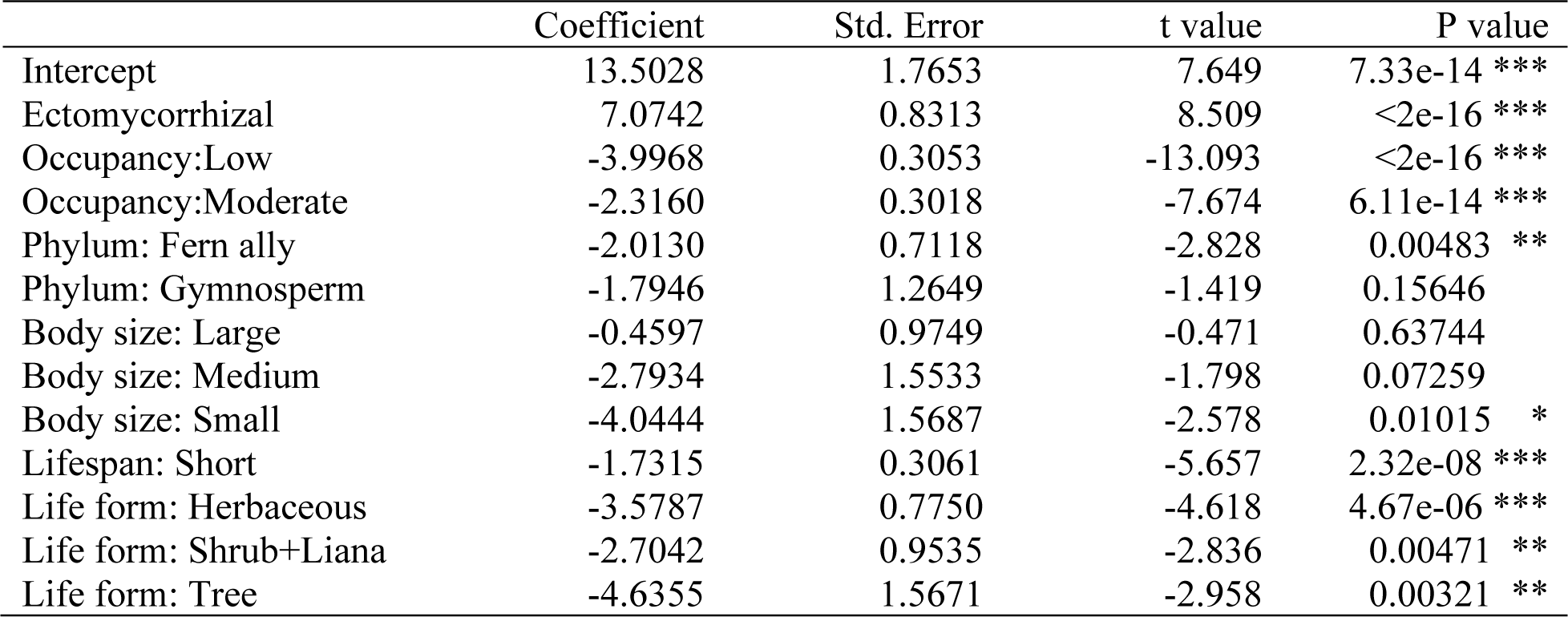
Regression model of arthropod link score per plant taxon as predicted from plant attributes. Overall model adjusted R-squared 0.4521; F-statistic 46.52 on 12 and 650 DF, p-value << 0.0001

|  | Coefficient | Std. Error | t value | P value |
| --- | --- | --- | --- | --- |
| Intercept | 13.5028 | 1.7653 | 7.649 | 7.33e-14 *** |
| Ectomycorrhizal | 7.0742 | 0.8313 | 8.509 | <2e-16 *** |
| Occupancy:Low | -3.9968 | 0.3053 | -13.093 | <2e-16 *** |
| Occupancy:Moderate | -2.3160 | 0.3018 | -7.674 | 6.11e-14 *** |
| Phylum: Fern ally | -2.0130 | 0.7118 | -2.828 | 0.00483 ** |
| Phylum: Gymnosperm | -1.7946 | 1.2649 | -1.419 | 0.15646 |
| Body size: Large | -0.4597 | 0.9749 | -0.471 | 0.63744 |
| Body size: Medium | -2.7934 | 1.5533 | -1.798 | 0.07259 |
| Body size: Small | -4.0444 | 1.5687 | -2.578 | 0.01015 * |
| Lifespan: Short | -1.7315 | 0.3061 | -5.657 | 2.32e-08 *** |
| Life form: Herbaceous | -3.5787 | 0.7750 | -4.618 | 4.67e-06 *** |
| Life form: Shrub+Liana | -2.7042 | 0.9535 | -2.836 | 0.00471 ** |
| Life form: Tree | -4.6355 | 1.5671 | -2.958 | 0.00321 ** |

The bivariate models showed in general that observed and predicted link sum were roughly equally good predictors, with the observed link sum working slightly better in most cases. Only for internally feeding phytophagous arthropods, the predicted link sum yielded much better than the observed link sum (Fig. 2).

**Fig. 2.**
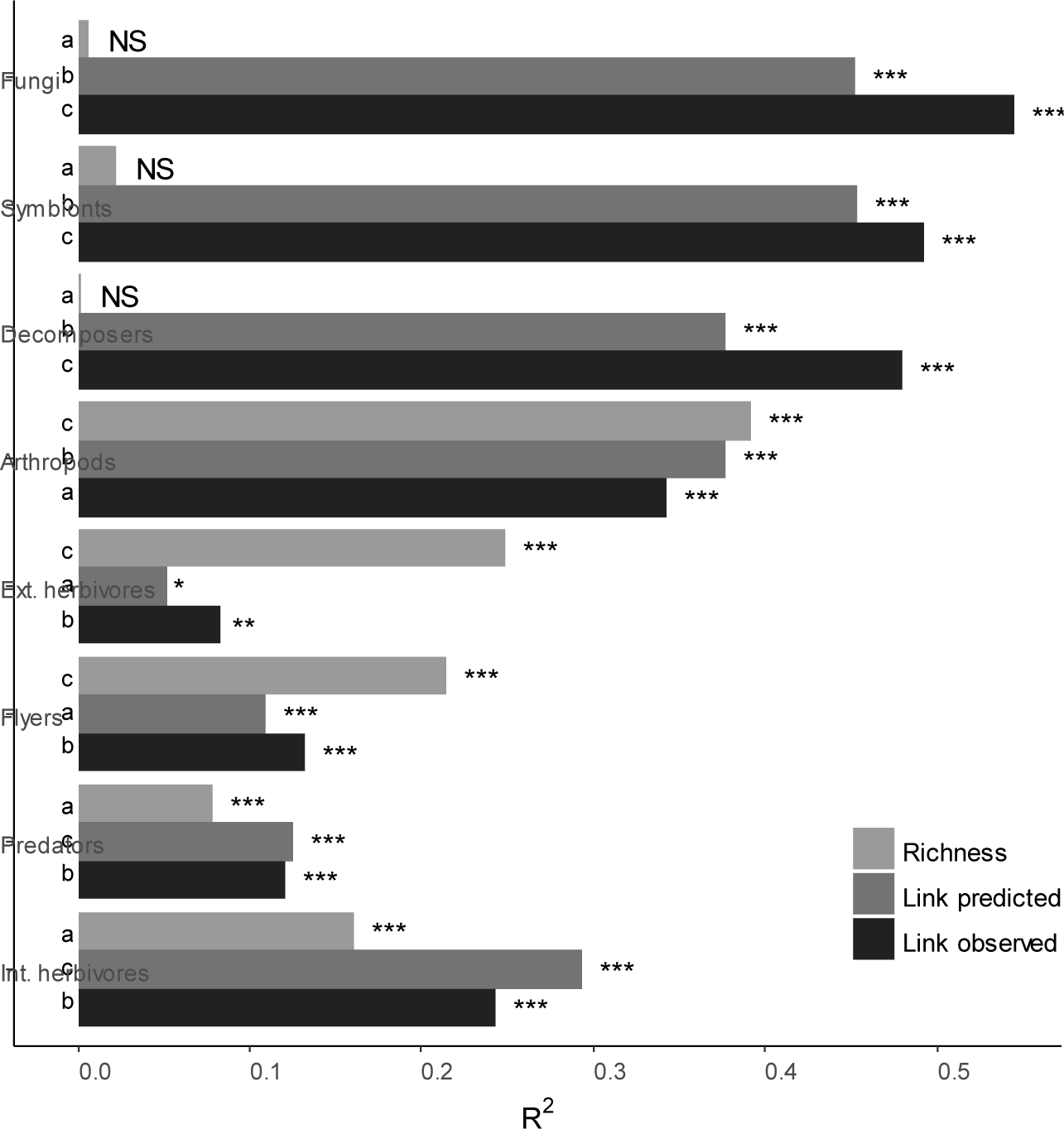
Bivariate models. Bars represent pseudo-R^2^ for regression models of consumer species richness in response to simple vascular plant species richness and plant richness weighted link scores for consumer associations. Significance of plant richness or link effects

Comparing the predictive power of link sum (link-weighted plant species richness) to simple plant richness gave somewhat contrasting results for fungi and arthropods. For fungal richness, simple plant species richness was a very poor predictor. In contrast, both observed and predicted link sum were quite strong and significant predictors of fungal richness, and with observed link sum providing consistently better modelling results than predicted link sum, accounting for more than 50% of the variation in fungal richness (Fig. 2). Moreover, this result was almost equally attributable to decomposer and symbiotic fungi. For genetic richness of soil fungi (OTU Fungi), in contrast, simple plant species richness outperformed link sum. The modelling outcome of the bivariate models was very different for total arthropod richness and most arthropod subgroups, for which plant richness was a superior predictor. However, for internally feeding phytophagous arthropods (gallers and miners), link sum performed markedly better than simple plant richness, while for predatory arthropods, only somewhat better. The best models reached 39% explained variation for total arthropod richness (simple plant richness) and 29% explained variation for internal feeders (predicted link score).

After fitting environmental proxies (community mean Ellenberg Indicator Values) as covariates in a multiple regression model, plant species richness became a significant predictor of all three measures of fungal richness. However, link sum, observed and predicted alike, remained stronger predictors of fungi richness and richness of most arthropod response groups than simple species richness. Observed link score was generally stronger than predicted link score, although the difference was modest. The multiple regression model for total fungi richness using observed link score reached 55% explained variation. Multiple regression including environmental proxies only contributed with a minor improvement of predictions compared to the best bivariate model (54.5% explained variation). Environmental proxies alone explained less than half of the variation of fungi richness, a little more than half of the variation for decomposers and one third of the variation in fungal symbiont richness (Fig. 3).

**Fig. 3.**
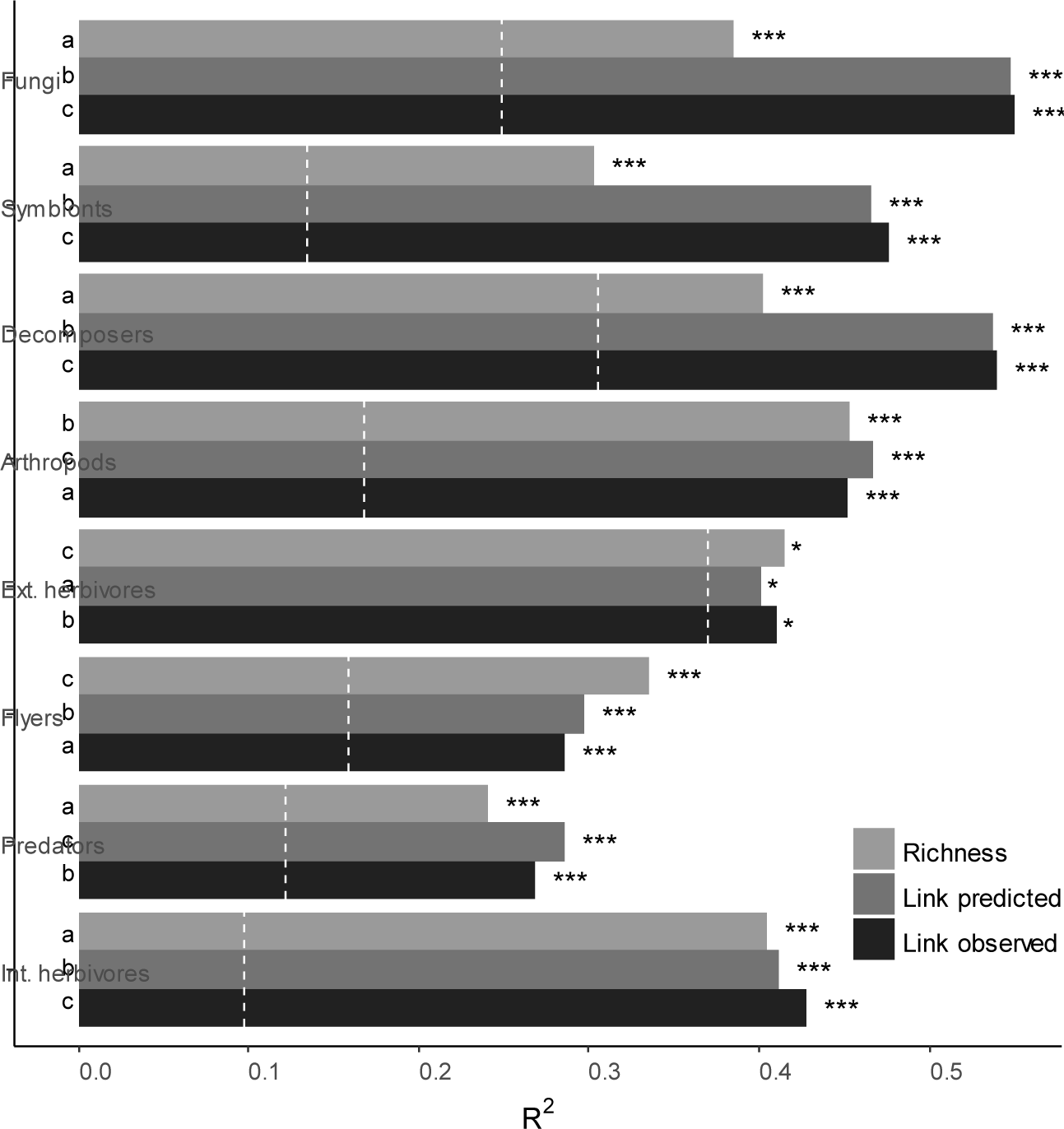
Multiple regression. Bars represent pseudo-R^2^ for regression models of consumer species richness in response to simple vascular plant species richness and plant richness weighted link scores for consumer associations, and with environmental calibration by mean Ellenberg Indicator Values as co-variables. The R^2^ for a model on environmental calibration alone is indicated by white lines.

Arthropod models improved markedly after fitting Ellenberg variables, with 47% explained variation for total arthropod richness (predicted link sum), 43% for internal feeders (observed link sum) and 42% for external feeders (simple plant richness). The difference between performance of plant richness and link scores was reduced compared to the bivariate models. The amount of variation explained by environmental proxies alone was low for internal feeders and total arthropod richness (23 and 36%, respectively), moderate for predators and flyers (43% and 47%, respectively) and high for external feeders (89%).

## Discussion

An overarching question of this study was whether plant species community composition may work as a predictor of consumer species richness. In a previous study in the same system of 130 sites, we showed that plants predict general species richness across taxonomic and functional groups, given that predictions are based on plant species richness amended with plant-based bioindication of habitat properties (Brunbjerg *et al.* 2018). In the present study, we have demonstrated that plant surrogacy of multi-taxon biodiversity can be taken a step further by including the value to consumer biodiversity of individual plant species.

The coterie of plant-associated arthropods and fungi varies considerably and predictably across plant taxa, a notion that was firmly established more than half a century ago (Southwood 1961). The results obtained in the present analysis are, for the most part, well aligned with the rich body of literature on the topic. Large-bodied, long-lived, structurally complex, widely distributed and locally abundant plant species have repeatedly been shown to harbour a larger fauna of phytophagous arthropods (e.g. Tahvanainen & Niemelä 1987; Brändle & Brandl 2001) and fungi (Strong & Levin 1975; Miller 2012). This has been encapsulated in the concept of apparency, which comprises both species’ attributes (e.g. body size and longevity) and of their history of immigration and fate in community dynamics (regional occupancy and local abundance). We found plant taxon nativeness, which is positively related to time since immigration, to be a correlate of consumer richness, but only in the model for fungi richness and, interestingly, only nativeness on the scale of Europe, not Denmark. Thus, it seems that associated consumers effectively track their host plants on the larger regional scale.

The capacity of plant taxa to form ectomycorrhiza (ECM) was little surprising as a predictor for associated fungal richness. It was, however, not anticipated that ectomycorrhizal capacity was a strong predictor of associated arthropod richness. This surprising pattern was not simply due to the fact that almost all ectomycorrhizal host plants are trees, as plant body size and growth form were also included as model predictors. Thus, within life-form groups, ectomycorrhizal plant taxa are on average hosts to a larger arthropod fauna than are non-ECM taxa. For trees and shrubs, genera such as *Fagus, Quercus, Betula* and *Salix* host more arthropod species than do *Ulmus, Acer, Fraxinus* and *Crataegus*, and similarly the dwarf-shrub *Salix repens* has a richer associated fauna than other similar-sized plant species. The mechanism behind this non-random co-occurrence escapes explanation, but calls for more detailed investigations.

When predicting observed species richness of arthropods and fungi in actual communities, models based on simple plant traits (predicted link sum) generally performed almost as good as models based on databased interaction links (observed link sum) or even better for all pooled arthropods. This result is encouraging for the use of plants in biodiversity surrogacy outside the study region used here. Basic knowledge on fungas and arthropod faunas is very far from complete in large parts of the world, and much more so than vascular floras (Mora *et al.* 2011; Hawksworth & Lücking 2017), and knowledge on species’ host relations is even more incomplete. In contrast, simple plant traits, such as life form and body size, are available for almost all plant species on the Globe, at least on a coarse scale. Thus, our finding is promising, and calls for further validation across global biomes.

The predictive power of interaction link sum on observed consumer richness varied considerably across functional groups of fungi and arthropods (Fig. 2). The effect of plant properties was strongest on the richness of biotrophs, decomposers and internally feeding arthropods. To a large extent, the physical and chemical properties of host plants define the habitat of species in these groups, which live in close intimacy with their host (Mazziotta *et al.* 2017) and cannot escape factors such as chemical plant defence, not even as after-life effects (Purahong *et al.* 2018). The small effect for externally feeding arthropods was surprising at first glance, because these species are herbivores and quite many of them oligophagous. On the other hand, many externally feeding phytophagous insects are associated with habitat type, such as lake margins or heathlands, and use a variety of host plants within that habitat, e.g. taxa such as the leaf beetles (Chrysomelidae) and the plant bugs (Miridae). The richness of external feeders, thus, was largely predictable from general habitat conditions derived from plant community composition through bioindication (Fig. 3). For predators, in contrast, one could think that “meat is meat” and plant species identity would have no effect. Nonetheless, we found secondary consumer richness to have a direct relationship with plant species richness. This communication between the first and the third trophic levels may be because many predators, in particular insect parasitoids, are quite host specific and use plant species chemistry as cue to locate their host (Godfray 1995).

There was an appreciable indirect effect of extrinsic habitat conditions – *position* in ecospace (Brunbjerg *et al.* 2017b) – on the observed consumer richness (Fig. 3), yet we could clearly demonstrate an added effect of interaction link sum across all functional groups. This effect was particularly evident for fungi, both biotrophs and decomposers. For arthropods, the additive predictive power of link sums over simple plant richness was generally dwarfed. However, a strong effect of plant richness remained on top of environmental calibration and after including environmental co-variates, predicted link score turned out to be the most significant predictor.

A core component in the *ecospace* approach to understanding consumer biodiversity is the diversification of carbon pools. While there is no easy way to directly characterize and classify different pools of dead organic matter in ecosystems, the classification of plants offer an opportunity for investigating the importance of carbon diversification for heterotrophic diversity. However, important carbon pools such as dung, carcass and dead wood, which were not part of our assessment, deserve further investigations.

Our results lend support to the notion that site-level biodiversity is an emergent property of site conditions (Brunbjerg et al. 2020 OIKOS), within the bounds of the regional species pool. Likewise, the results demonstrate that biodiversity begets biodiversity, with community-level plant species richness in the role as a central bottom-up driver with strong effects across taxonomic groups, trophic levels and the parasitic-mutualistic-saprotrophic continu<u>um</u>a (Põlme *et al.* 2018). Our results may be applied in conservation science in order to improve the evaluation of planning and management choices, also in areas without much knowledge of the consumer biotas and their host relationships. Further, our results may be applied to novel ecosystem in the management of urban biodiversity.

## Author contributions statement

HHB, RE, AKB and JHC conceived the ideas and designed the methodology; AKB, IG, TL, SH, HHB, RE, TGF, JHC and CF collected the data; RE, AKB, HHB and LD analysed the data; HHB, RE and AKB led the writing of the manuscript. All authors contributed critically to the drafts and gave final approval for publication.

## Acknowledgements

RE, TGF, TL and AKB were supported by a grant from VILLUM foundation (Biowide, VKR-023343). We thank all volunteers that have helped in data collection, Karl-Henrik Larsson for aid in identifying critical corticioid fungi, Leif Örstadius for identifying Psathyrella collections.

## References

Bocci, G. (2015) TR8: an R package for easily retrieving plant species traits. Methods in Ecology and Evolution, 6, 347–350.

Brunbjerg, A.K., Bruun, H.H., Broendum, L., Classen, A.T., Fog, K., Froeslev, T.G., Goldberg, I., Hansen, M.D.D., Hoeye, T.T., Laessoee, T., Newman, G., Skipper, L., Soechting, U. & Ejrnaes, R. (2017a) A systematic survey of regional multitaxon biodiversity: evaluating strategies and coverage. bioRxiv.

Brunbjerg, A.K., Bruun, H.H., Dalby, L., Fløjgaard, C., Frøslev, T.G., Høye, T.T., Goldberg, I., Læssøe, T., Hansen, M.D.D., Brøndum, L., Skipper, L., Fog, K. & Ejrnæs, R. (2018) Vascular plant species richness and bioindication predict multi-taxon species richness. Methods in Ecology and Evolution, 9, 2372–2382.

Brunbjerg, A.K., Bruun, H.H., Moeslund, J.E., Sadler, J.P., Svenning, J.-C. & Ejrnæs, R. (2017b) Ecospace: A unified framework for understanding variation in terrestrial biodiversity. Basic and Applied Ecology, 18, 86–94.

Brändle, M. & Brandl, R. (2001) Species richness of insects and mites on trees: expanding Southwood. Journal of Animal Ecology, 70, 491–504.

Brändle, M. & Brandl, R. (2003) Species richness on trees: a comparison of parasitic fungi and insects. Evolutionary Ecology Research, 5, 941–952.

Brändle, M. & Brandl, R. (2006) Is the composition of phytophagous insects and parasitic fungi among trees predictable? Oikos, 113, 296–304.

Buchwald, E., Wind, P., Bruun, H.H., Møller, P.F., Ejrnæs, R. & Svart, H.E. (2013) Hvilke planter er hjemmehørende i Danmark? Flora og Fauna, 118, 73–96.

Ellenberg, H., Weber, H.E., Düll, R., Wirth, V., Werner, W. & Paulissen, D. (1991) Zeigerwerte von Pflanzen in Mitteleuropa.

Verlag E. Goltze KG, Göttingen. Feeny, P. (1976) Plant apparency and chemical defense. Biological Interactions Between Plants and Insects (eds J.W. Wallace & R.L. Nansel), pp. 1–40. Plenum Press, New York, USA.

Godfray, H.C.J. (1995) Communication between the first and third trophic levels: An analysis using biological signalling theory. Oikos, 72, 367–374.

Hansen, K. (2004) Dansk Feltflora, 1. udg. 9. opl. edn. Gyldendal, Copenhagen.

Hartvig, P. & Vestergaard, P. (2015) Atlas Flora Danica. pp. 1230. Gyldendal, København.

Hawksworth, D.L. (2001) The magnitude of fungal diversity: the 1.5 million species estimate revisited. Mycological Research, 105, 1422–1432.

Hawksworth, D.L. & Lücking, R. (2017) Fungal diversity revisited: 2.2 to 3.8 million species. Microbiology Spectrum, 5.

Heilmann-Clausen, J., Maruyama, P.K., Bruun, H.H., Dimitrov, D., Læssøe, T., Frøslev, T.G. & Dalsgaard, B. (2016) Citizen science data reveal ecological, historical and evolutionary factors shaping interactions between woody hosts and wood-inhabiting fungi. New Phytologist, 212, 1072–1082.

Hempel, S., Götzenberger, L., Kühn, I., Michalski, S.G., Rillig, M.C., Zobel, M. & Moora, M. (2013) Mycorrhizas in the Central European flora: relationships with plant life history traits and ecology. Ecology, 94, 1389–1399.

Kelly, C.K. & Southwood, T.R.E. (1999) Species richness and resource availability: a phylogenetic analysis of insects associated with trees. Proceedings of the National Academy of Sciences of the United States of America, 96, 8013–8016.

Kemp, J.E. & Ellis, A.G. (2017) Significant local-scale plant-insect species richness relationship independent of abiotic effects in the temperate Cape floristic region biodiversity hotspot. PLoS ONE, 12, e0168033.

Kennedy, C.E.J. & Southwood, T.R.E. (1984) The number of species of insects associated with British trees: a re-analysis. Journal of Animal Ecology, 53, 455–478.

Kleyer, M., Bekker, R.M., Knevel, I.C., Bakker, J.P., Thompson, K., Sonnenschein, M., Poschlod, P., van Groenendael, J.M., KlimeŠ, L., Klime Š ová, J., Klotz, S., Rusch, G.M., Hermy, M., Adriaens, D., Boedeltje, G., Bossuyt, B., Dannemann, A., Endels, P., Götzenberger, L., Hodgson, J.G., Jackel, A.K., Kühn, I., Kunzmann, D., Ozinga, W.A., Römermann, C., Stadler, M., Schlegelmilch, J., Steendam, H.J., Tackenberg, O., Wilmann, B., Cornelissen, J.H.C., Eriksson, O., Garnier, E. & Peco, B. (2008) The LEDA Traitbase: a database of life-history traits of the Northwest European flora. Journal of Ecology, 96, 1266–1274.

Klotz, S., Kühn, I. & Durka, W. (2002) BIOLFLOR – Eine Datenbank mit biologisch-ökologischen Merkmalen zur Flora von Deutschland. Bundesamt für Naturschutz / Landwirtschaftsverlag, Bonn.

Larsen, F.W., Bladt, J. & Rahbek, C. (2009) Indicator taxa revisited: useful for conservation planning? Diversity and Distributions, 15, 70–79.

Lawton, J.H. & Schröder, D. (1977) Effects of plant type, size of geographical range and taxonomic isolation on number of insect species associated with British plants. Nature, 265, 137–140.

Lindenmayer, D., Barton, P., Westgate, M., Lane, P. & Pierson, J. (2015) Biodiversity surrogates. Indicators and Surrogates of Biodiversity and Environmental Change (eds D. Lindenmayer, P. Barton & J. Pierson), pp. 15–24. CRC Press, Boca Raton, Florida, USA.

Mazziotta, A., Vizentin-Bugoni, J., Tøttrup, A.P., Bruun, H.H., Fritz, Ö. & Heilmann-Clausen, J. (2017) Interaction type and intimacy structure networks between forest-dwelling organisms and their host trees. Basic and Applied Ecology, 24, 86–97.

Miller, Z.J. (2012) Fungal pathogen species richness: why do some plant species have more pathogens than others? The American Naturalist, 179, 282–292.

Mora, C., Tittensor, D.P., Adl, S., Simpson, A.G.B. & Worm, B. (2011) How many species are there on Earth and in the Ocean? PLoS Biology, 9, e1001127.

Nilsson, R.H., Larsson, K.-H., Taylor, A.F.S., Bengtsson-Palme, J., Jeppesen, T.S., Schigel, D., Kennedy, P., Picard, K., Glöckner, F.O., Tedersoo, L., Saar, I., Kõljalg, U. & Abarenkov, K. (2019) The UNITE database for molecular identification of fungi: handling dark taxa and parallel taxonomic classifications. Nucleic Acids Research, 47, D259–D264.

Põlme, S., Bahram, M., Jacquemyn, H., Kennedy, P., Kohout, P., Moora, M., Oja, J., Öpik, M., Pecoraro, L. & Tedersoo, L. (2018) Host preference and network properties in biotrophic plant–fungal associations. New Phytologist, 217, 1230–1239.

Purahong, W., Wubet, T., Krüger, D. & Buscot, F. (2018) Molecular evidence strongly supports deadwood-inhabiting fungi exhibiting unexpected tree species preferences in temperate forests. The ISME Journal, 12, 289–295.

Savile, D.B.O. (1979) Fungi as aids to plant taxonomy: methodology and principles. Symbolae Botanicae Upsalienses, 22, 135–145.

Seibold, S., Bässler, C., Brandl, R., Büche, B., Szallies, A., Thorn, S., Ulyshen, M.D. & Müller, J. (2016) Microclimate and habitat heterogeneity as the major drivers of beetle diversity in dead wood. Journal of Applied Ecology, 53, 934–943.

Southwood, T.R.E. (1961) The number of species of insect associated with various trees. Journal of Animal Ecology, 30, 1–8.

Strong, D.R., Lawton, J.H. & Southwood, T.R.E. (1984) Insects on Plants: Community Patterns and Mechanisms. Blackwell Scientific Publications, Oxford.

Strong, D.R. & Levin, D.A. (1975) Species richness of parasitic fungi of British trees. Proceedings of the National Academy of Sciences of the United States of America, 72, 2116–2119.

Strong, D.R. & Levin, D.A. (1979) Species richness of plant parasites and growth form of their host. American Naturalist, 114, 1–22.

Tahvanainen, J. & Niemelä, P. (1987) Biogeographical and evolutionary aspects of insect herbivory. Annales Zoologici Fennici, 24, 239–247.

Tedersoo, L., Bahram, M., Cajthaml, T., Põlme, S., Hiiesalu, I., Anslan, S., Harend, H., Buegger, F., Pritsch, K., Koricheva, J. & Abarenkov, K. (2015) Tree diversity and species identity effects on soil fungi, protists and animals are context dependent. ISME J, 10, 346–362.

